# A high-throughput screening dataset of small-molecule inhibitors across human DNA glycosylases

**DOI:** 10.64898/2026.09.25.752594

**Authors:** Alice Eddershaw, Bjørn Dalhus, Opher Gileadi, Susanne Gräslund, İrşil Güneş, Thomas Helleday, Kang-Cheng Liu, Olga Loseva, Vu To Nakstad, Hilde Loge Nilsen, Nicola P. Montaldo, Karen Nierlin, Carina Norström, Natálie Rudolfová, Kristine Sletta, Michael Sundström, Rahul Upadhyay, Barbara van Loon, Edvard Wigren, Elisée Wiita, Ane Marit Wågbø, Kaixin Zhou, Evert J. Homan, Torkild Visnes, Maurice Michel

## Abstract

The base excision repair pathway removes small base lesions from DNA and is initiated in humans by one of eleven DNA glycosylases with overlapping substrate specificities. Despite their central role in genome maintenance and transcription, systematic datasets describing small-molecule binders or inhibitors of DNA glycosylases are lacking. Here, through a collaborative effort across Scandinavia, we establish a high-throughput biochemical assay platform for human DNA glycosylases and use it to generate a systematic small-molecule screening dataset. A chemogenomic library was screened against nine glycosylases with assays of sufficient quality for hit identification, yielding primary screening and concentration-response data and identifying multiple previously unreported inhibitors. All primary and processed data, along with detailed assay protocols and metadata, are publicly available to support reuse in chemical biology, further compound optimization, assay development and comparative studies of DNA repair enzymes.

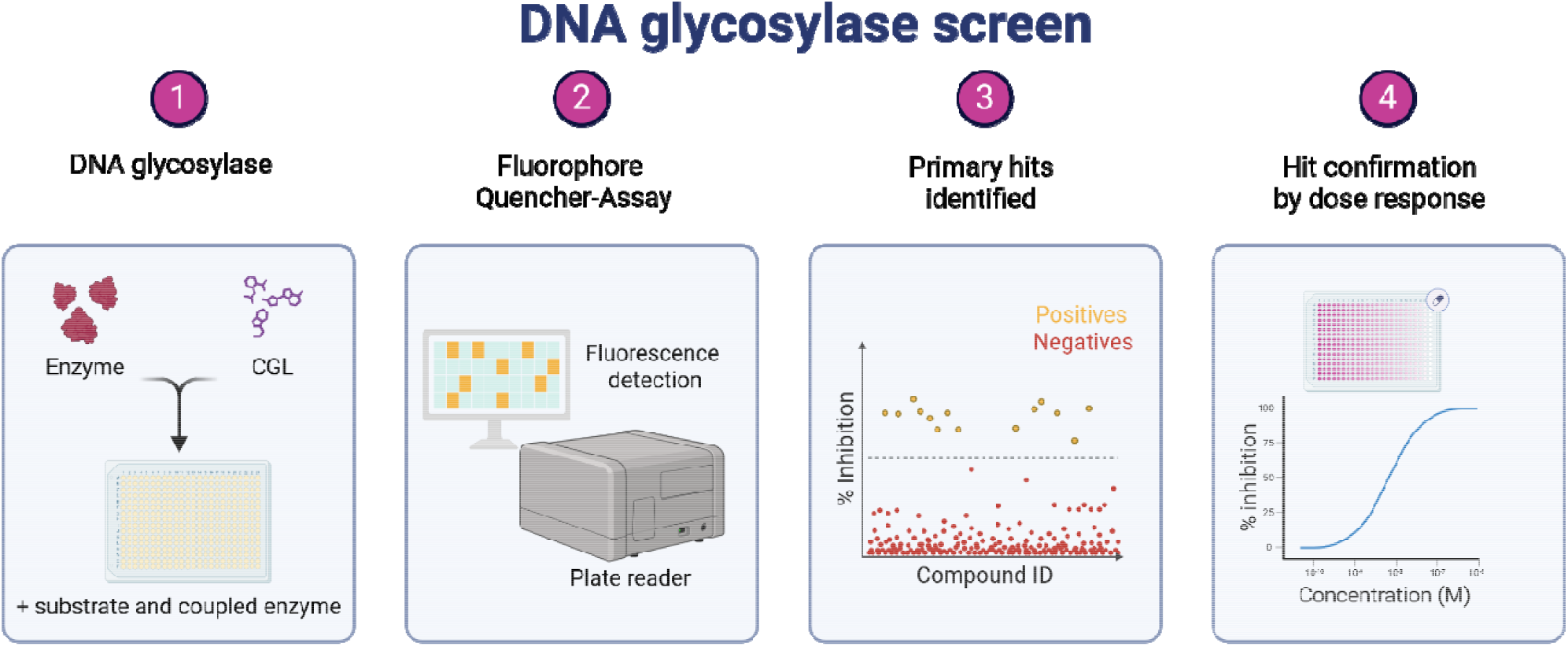

## Introduction

The base excision repair (BER) pathway is a central mechanism for the removal of small base lesions from DNA, including oxidation, alkylation, mismatch and deamination products.^1^ In humans, BER is initiated by one of eleven DNA glycosylases that recognize structurally diverse but partially overlapping substrates and excise damaged or mispaired bases from DNA. Several members of this family are bifunctional, additionally possessing apurinic/apyrimidinic (AP) lyase activity enabling incision of the DNA backbone, whereas others function solely as monofunctional glycosylases.

Despite extensive biochemical and structural characterization, systematic chemical tools for probing DNA glycosylase function remain limited.^2-4^ Most reported small-molecule studies have focused on individual enzymes,^5-7^ and comprehensive datasets describing small-molecule binding broadly across the human DNA glycosylases are notably rare.^8^ The absence of such resources has limited analysis of ligand recognition, assay development efforts and the broader exploration of chemical modulation within these enzymes.^9,10^

Here, we coordinate and collaborate across multiple laboratories in Norway and Sweden and assemble nine human DNA glycosylases in one modular high-throughput format, enabling rapid selectivity and potency assessment. Using this assay platform based on quencher-fluorophore DNA constructs, a subset of a recently developed chemogenomic library (CGL) of small molecules was screened against each enzyme, generating quantitative primary and validation measurements of small-molecule inhibition and binding. The dataset includes assay performance metrics, control compounds, concentration-response data for selected hits, associated metadata and validation by thermal stabilisation. Importantly, the CGL contains data of overlapping chemotypes and analogues, informing molecular patterns for individual enzyme targets. All data and protocols are made publicly available to support reuse in chemical biology, DNA repair research and the development of new assays and, finally, selective chemical probes targeting DNA glycosylases.^11^

This dataset expands the collection of reported inhibitors for multiple DNA glycosylases and provides, for several enzymes, the first publicly available examples of small-molecule engagement. The resulting resource enables comparative analyses of target ligandability and provides starting points for future medicinal chemistry campaigns.

## Methods

### Assay principle

DNA glycosylase activity was measured using a fluorescence-based DNA strand incision assay as reported before.^8,12-14^ Synthetic double-stranded DNA substrates contained a site-specific damaged or modified base recognized by the respective DNA glycosylase, along with a 5′-fluorophore (6-carboxyfluorescein, FAM, or Sulfo-Cyanine 5, Cy5). A quencher (Dimethyl-aminoazo-benzolsulfonyl, Dabcyl, or Black Hole Quencher 2, BHQ2) was incorporated on the 3′ complementary strand opposite the fluorophore. In the intact duplex, fluorescence emission is suppressed due to proximity of the quencher. Upon excision of the damaged base and subsequent strand incision, separation of fluorophore and quencher results in increased fluorescence signal. For monofunctional DNA glycosylases, including AAG (AAG), MUTYH, SMUG1, UNG, MBD4, TDG and OGG1, strand incision was enabled by coupling base excision to apurinic/apyrimidinic endonuclease 1 (APE1). Bifunctional DNA glycosylases possessing intrinsic AP-lyase activity, including NEIL1, NEIL2, NEIL3 and NTHL1, were assayed without APE1. Fluorescence was monitored continuously or at defined time points to quantify enzymatic turnover and compound-dependent modulation of activity.

### Recombinant Proteins

Proteins were expressed and purified as described here for NTHL1^15^, and for AAG, NEIL1, TDG, OGG1, APE1 here^8,14,16^. For MBD4, a pNIC28 expression vector encoding the full-length human gene with an N-terminal 6xHis tag and thrombin cleavage site was used. For SMUG1, a pET29a vector encoding a codon-optimised gene for wild-type hSMUG1 with a TEV-protease cleavable N-terminal 6xHis tags and C-terminal AviTag was synthesized by Twist BioScience. Vectors were transformed into BL21 pRARE *E*.*coli* for expression. A single colony was used to inoculate TB media with 50 µg/mL kanamycin and 35 µg/mL chloramphenicol and cells were grown overnight at 37 °C with 220 RPM. Starter culture was diluted 1:100 into fresh TB buffer with 50 µg/mL kanamycin and cells were grown to OD_600_ of 2 before inducing with 0.5 mM IPTG and culturing at 18 °C for 16 hours with 220 RPM. Cells were harvested and stored at -80 °C for purification. The cells were lysed in a buffer of 50 mM HEPES pH 7.5, 500 mM NaCl, 10 mM imidazole, 0.5 mM TCEP, 10% v/v glycerol and EDTA free-protease inhibitor tablets (Roche) by sonication for 5 minutes (5 seconds on, 25 seconds off) at 4 °C. DNA was precipitated by incubation with 0.2% PEI, and lysate clarified by 22,000 xg for 30 mins. The supernatant was filtered through a 0.22 um membrane and loaded onto a 5 mL HisTrap column using an ÄKTA system. The column was washed in 6 CV of lysis buffer with 30 mM imidazole, before eluting in lysis buffer with 300 mM imidazole. The peak fractions were pooled and concentrated using 30 kDa MWCO. The sample was loaded onto a S200 column for purification by size exclusion chromatography in a buffer of 10 mM HEPES pH 7.5, 150 mM NaCl, 5% glycerol and 0.5 mM TCEP. Fractions containing the target protein were assessed by SDS-PAGE and the pooled and concentrated stock of enzyme was stored in aliquots at -80 °C. The identify of the target protein was confirmed by LC-MS. NEIL3 was purified as reported before.^17^

### DNA substrates

Custom oligonucleotide substrates were synthesized with enzyme-specific lesions, including uracil, thymine glycol, 8-oxo-adenine, inosine or abasic site analogues and ordered from ATD Bio (Biotage) or IDTDNA (Table M1). Oligonucleotides containing 5’ Cy5 and 3’ BHQ2 were from GeneLink. Fluorophore- and quencher-labelled strands were annealed at a 1:1.25 molar ratio in annealing buffer (25 mM Tris-HCl pH 8.0, 50 mM NaCl, 2 mM MgCl_2_) by heating to 95°C followed by slow cooling to room temperature. Annealed substrates were stored at -20°C protected from light until use.

### DNA sequences

DNA sequences were designed to be complementary across the platform, i.*e*. having the same sequence except lesions and complementary base (Table M1). Sequences were as reported before ^18^: 5′-FAM-TCTG CCA <u>X</u>CA CTG CGT CGA CCT G-3′ and, complementary, 5′-CAG GTC GAC GCA GTG <u>Y</u>TG GCA GT-Dab-3 where X is the preferred lesion (8-oxoA, U, G, 5-OH-U) and Y the corresponding complementary base (T, G, C, A), Dab = Dabcylquencher, and FAM = 6-carboxyfluorescein. For NTHL1, NEIL1 and 2, Cy5 and BHQ2 labelled versions were used. For NEIL3, a single-stranded ‘bubble’ mimetic was used: 5′-FAM-TCTG CCA TGA AC<u>Tg</u> CGA GGC CAC TGC GTC GAC CT G-3′ and, complementary, 5′-CAG GTC GAC GCA GTG CTC CCT TGG AGC TGG CAG T-Dab-3′.

**Table M1:** Substrates for DNA glycosylase activity assays.

| Enzyme | Fluorophore | Quencher | Substrate Lesion | Complementary base | Context |
| --- | --- | --- | --- | --- | --- |
| AAG | FAM | Dabcyl | U | T | Duplex 23-mer DNA |
| MBD4 | FAM | Dabcyl | U | G | Duplex 23-mer DNA |
| MUTYH | FAM | Dabcyl | G | G | Duplex 23-mer DNA |
| NTHL1 | Cy5 | BHQ2 | 5-OH-U | A | Duplex 23-mer DNA |
| NEIL1 | Cy5 | BHQ2 | 5-OH-U | C | Duplex 23-mer DNA |
| NEIL2 | Cy5 | BHQ2 | 5-OH-U | C | Duplex 23-mer DNA |
| NEIL3 | FAM | Dabcyl | Tg | Single-stranded | Bubble-34 mer DNA |
| OGG1 | FAM | Dabcyl | 8oxoA | C | Duplex 23-mer DNA |
| SMUG1 | FAM | Dabcyl | U | G | Duplex 23-mer DNA |
| TDG | FAM | Dabcyl | U | G | Duplex 23-mer DNA |
| UNG | FAM | Dabcyl | U | G | Duplex 23-mer DNA |

### Enzymes and assay conditions

Recombinant human DNA glycosylases (OGG1, UNG2, TDG, SMUG1, AAG, NEIL1, NEIL2, NEIL3, MUTYH, MBD4, and NTHL1) were assayed under enzyme-specific buffer conditions optimized for activity and signal stability, using the EUbOPEN protocol for DNA glycosylases ^12^. Reaction buffers typically contained Tris-HCl (pH 8.0), NaCl or KCl, MgCl□ where required, and low concentrations of non-ionic detergent (Tween-20). For monofunctional glycosylases, reactions additionally contained APE1 at a final concentration of 2 nM to promote efficient strand incision.

Assays were performed in black 384-well microplates with a total reaction volume of 50 µL per well. Enzyme solutions were preincubated with compounds for 10 min at room temperature prior to substrate addition. Reactions were initiated by addition of DNA substrate and monitored by fluorescence detection (excitation 485 nm and emission 535 nm for FAM/Dabcyl, excitation 620 nm, emission 670 nm for Cy5/BHQ2) using a multimode plate reader. Reaction times varied depending on enzyme turnover characteristics and ranged from kinetic measurements at 0, 8, 15 and 30 min to extended incubations for low-turnover.

### Chemogenomic Library

Through the EUbOPEN consortium ^19^ we have generated a chemogenomic library (CGL) covering small molecules targeting a large part of the human proteome.^20^ Compounds included in the CGL were selected to be free from licensing restrictions, commercially available through standard vendors, and to provide coverage of multiple chemotypes for each target. Compared with chemical probes, CGL compounds generally exhibit a lower degree of target selectivity, while still offering useful target engagement across a broad range of biological pathways. Compound selection was informed by resources such asCHEMBLdb, Probes&Drugs and the IUPHAR/BPS Guide to Pharmacology.^21-25^All compounds within the CGL have undergone quality control by LC-MS and have been assessed for toxicity in two cell lines.

### Compound handling and screening format

Compounds were transferred to assay plates using acoustic dispensing or liquid handling systems to achieve the desired final concentrations while maintaining constant DMSO levels. Screening plates included positive control compounds where available, negative (vehicle-only) controls, and no-enzyme controls to monitor background fluorescence. Control wells were distributed across each plate to enable plate-wise normalization and quality assessment. Compounds were screened at 10 µM or at 1 µM where indicated.

### Counter-screening for APE1 activity

To identify compounds that interfered with APE1 rather than DNA glycosylase activity, a dedicated APE1 counter-screen was performed using an abasic-site DNA substrate selective for AP-endonuclease activity. Assay conditions and detection settings were analogous to those used for glycosylase assays, enabling direct comparison and exclusion of nonspecific APE1 inhibitors from analysis.

### Data processing and quality control

Raw fluorescence data were normalized to plate controls and expressed as percent inhibition relative to positive and negative controls. Assay quality was assessed using signal-to-background ratios and Z′ factors calculated on a per-plate basis. All raw and processed data, along with assay metadata and analysis templates, are provided in the associated data repository to support reuse and independent reanalysis.

### Differential scanning fluorimetry (nanoDSF)

Thermal stability measurements were performed using nano differential scanning fluorimetry (nanoDSF) to assess compound-dependent effects on protein unfolding. Recombinant enzymes were incubated at 5 µM in assay buffer with selected small-molecule compounds or DMSO vehicle control prior to thermal scanning. Measurements were carried out using a nanoTemper Prometheus instrument with samples loaded into quartz capillaries, with two technical replicates (capillaries) measured per condition. Temperature was increased by 1 °C /min from 20 to 90 °C, and protein unfolding was monitored by intrinsic tryptophan fluorescence as a function of temperature. Melting temperatures (T_m_) were determined from changes in the fluorescence signal and its first derivative.

## Results

### Assay development and optimization for DNA glycosylases

To establish robust and comparable screening conditions across the human DNA glycosylases, enzyme titration and time-resolved activity measurements were performed for each enzyme prior to primary screening. For each DNA glycosylase, increasing concentrations of enzyme were incubated with a fixed concentration (10 nM) of the corresponding fluorescent DNA substrate, and fluorescence signal was monitored over time to assess reaction kinetics and signal development.

Kinetic readouts revealed enzyme concentration-dependent increases in fluorescence signal, with higher enzyme concentrations producing faster signal onset and earlier plateau formation. Lower enzyme concentrations resulted in slower reaction kinetics and reduced signal amplitudes, enabling identification of conditions under which product formation was linear over the selected assay window. Lower enzyme concentrations plateaued for selected members, indicating single turnover reactions of the enzymes on substrates. Representative enzyme titration and kinetic traces for nine DNA glycosylases and APE1 are shown in Figure 1.

**Figure 1.**
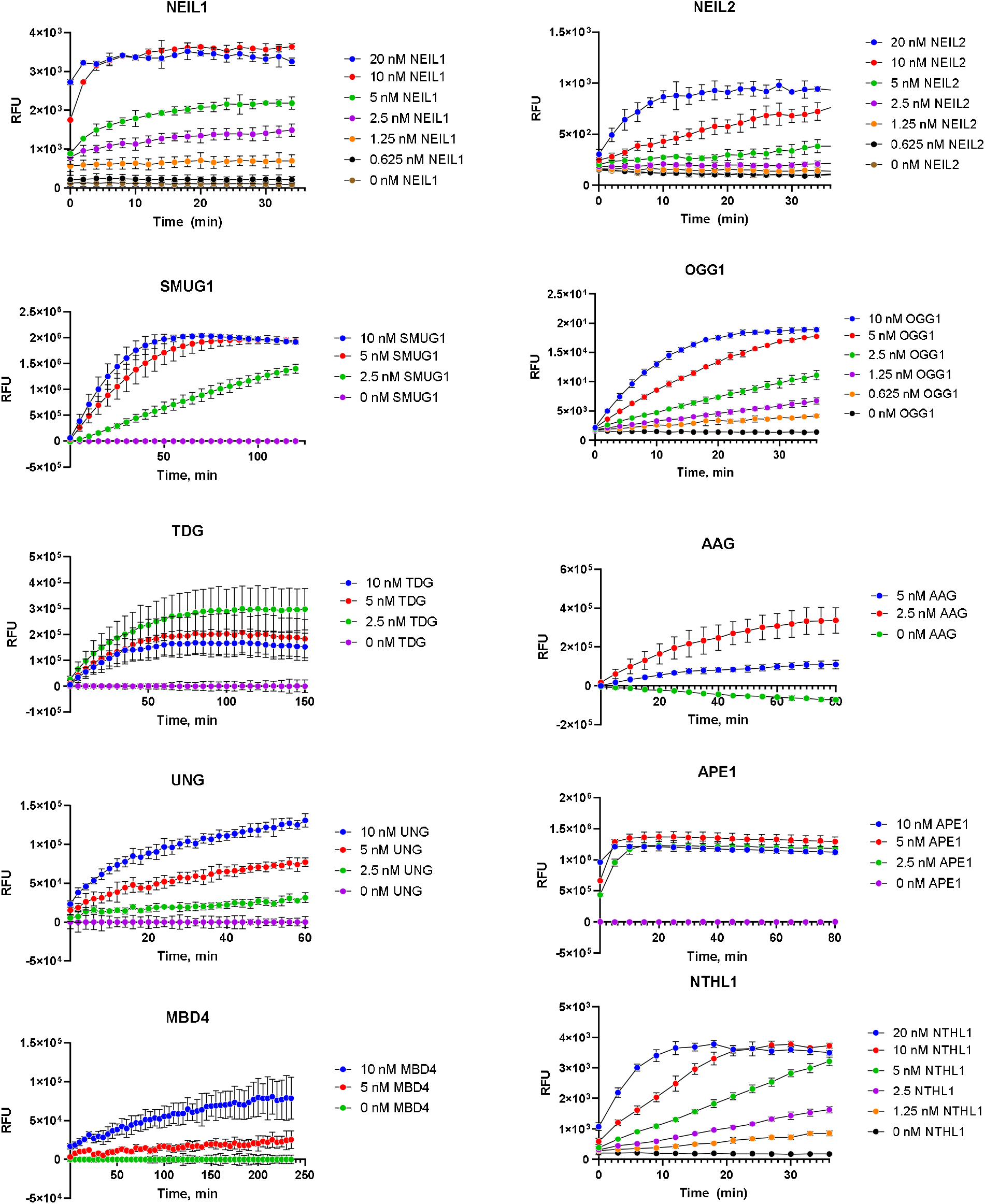
Concentration-dependent kinetics of DNA glycosylase enzymes measured by fluorescence assay. Relative fluorescence units (RFU) were monitored over time following incubation of substrate DNA with increasing concentrations of the indicated repair enzymes. Left panel: NEIL1, SMUG1, TDG, UNG2, and MBD4. Right panel: NEIL2, OGG1, AAG (AAG), APE1, and NTHL1. Error bars indicate mean ± SD. Note: The proteins were measured across three locations with as many setups. Variability in the readout resulted.

Based on these titration experiments, enzyme concentrations were selected to balance assay sensitivity, signal stability and throughput suitability. Final assay conditions were chosen to ensure sufficient dynamic range within the measurement window while minimizing substrate depletion and background signal. Enzyme concentrations used for primary screening and validation assays were derived directly from these optimization experiments and were determined individually for each DNA glycosylase – OGG1 (10 nM), AAG (10 nM), MBD4 (10 nM), UNG (10 nM), TDG (10 nM), SMUG1 (10 nM), MUTYH (10 nM), NEIL1 (4 nM), NEIL2 (4 nM), NEIL3 (10 nM), NTHL1 (2 nM). Equivalent titration and kinetic optimization experiments were performed for all eleven human DNA glycosylases included in the screening platform, and the corresponding data are provided above and in the supplementary materials. These assay-development datasets support the reproducibility and comparability of the screening results across the enzyme family.

### Primary screening of the CGL against all DNA glycosylases

Following assay optimisation and assessment of assay performance, nine DNA glycosylases were taken forward for primary screening analysis. MUTYH and NEIL3 were excluded because the respective assays did not meet the predefined quality criteria required for reliable hit identification. A subset of the EUbOPEN CGL, comprising 980 unique compounds was screened against the remaining DNA glycosylase panel. Enzyme activity was normalized to the activity in DMSO and no-enzyme controls, and inhibition thresholds set on a plate-specific basis (>3xSD of controls). Primary screening identified 70 compound–target hits of which 43 compounds were unique, with target-specific hit rates ranging from 0.2% to 2.1% (Table 1, Figure 2, Figure S1). The number of hits varied considerably between individual glycosylases. OGG1 showed the highest hit rate, with 21 compounds identified (2.1%), followed by TDG with 12 hits (1.2%), and MBD4 and NEIL2 with 10 hits each (1.0%). Lower hit rates were observed for the remaining enzymes, with between two and five compounds identified per target. Overall, the relatively low hit rates indicate that activity within the CGL was restricted to a small subset of compounds rather than reflecting widespread inhibition across the panel.

**Table 1.** Summary of primary screening and confirmed hits across the DNA glycosylase panel. The CGL comprising 980 unique compounds was screened against the indicated DNA glycosylases. The number of primary hits and corresponding hit rate are shown for each target, together with the number of inhibitors confirmed by concentration-response analysis. Hit rates were calculated relative to the total number of compounds screened.

| TARGET | # HITS | HIT RATE % | # CONFIRMED HITS |
| --- | --- | --- | --- |
| **MBD4** | 10 | 1.0 | 1 |
| **AAG** | 5 | 0.5 | 3 |
| **NEIL1** | 2 | 0.2 | 1 |
| **NEIL2** | 10 | 1.0 | 1 |
| **NTHL1** | 5 | 0.5 | 3 |
| **OGG1** | 21 | 2.1 | 18 |
| **SMUG1** | 3 | 0.3 | 1 |
| **TDG** | 12 | 1.2 | 6 |
| **UNG** | 2 | 0.2 | 2 |

**Figure 2.**
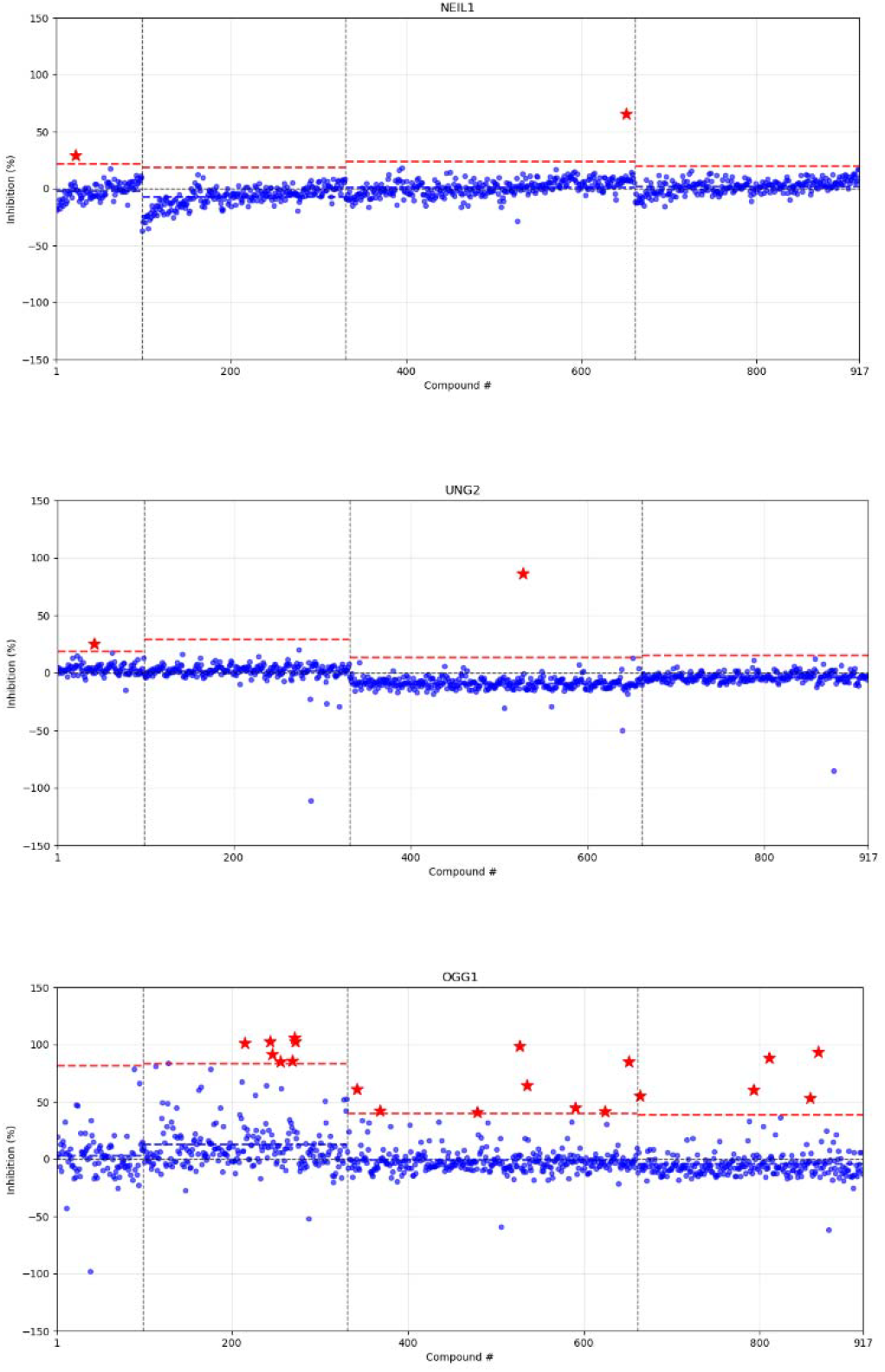
Primary screening of the CGL against human DNA glycosylases NEIL1, UNG and OGG1. Compounds were screened against the indicated DNA glycosylases and activity is shown as percent inhibition relative to plate controls. Each blue point represents an individual compound and red stars indicate compounds selected as primary hits. Red dashed lines indicate the plate-specific thresholds used for hit selection, and vertical dashed lines separate individual screening plates. Compound numbers correspond to their position within the screening library. Remaining members are shown in Figure S1.

Primary hits were subsequently prioritised for concentration-response analysis based on the magnitude and reproducibility of inhibition. Further compound availability and chemical quality, if applicable were considered. In total, 63 compounds were selected for follow-up across the panel, of which 37 were confirmed as inhibitors in concentration-response experiments, corresponding to an overall confirmation rate of 58.7%. The number of primary hits, compounds taken forward and confirmed inhibitors for each DNA glycosylase is summarised in Table 1.

### Hit validation

Hits identified in the primary screen were evaluated in dose-response assays alongside positive control compounds reported previously.^8,26,27^ As an AP site stalling compound, TH8777 was used as the positive control for MBD4, AAG, NTHL1, NEIL1, and NEIL2; TH10253 is a specific and potent inhibitor for OGG1; TH16142 and TH15932 are unpublished internal inhibitors for UNG and SMUG1 respectively. Of the 63 compounds selected for follow-up, 37 were confirmed as inhibitors in concentration-response experiments, corresponding to a validation rate of 58.7%. This relatively high confirmation rate supports the robustness of the primary screening workflow and indicates that the majority of selected hits represented reproducible enzyme-modulating compounds. Potencies span approximately one order of magnitude and pIC_50_ values range from 4.2 to 6.0 (corresponding to IC_50_ values from approximately 1 µM to 100 µM), as summarised in Figure 3 and in Tables S1. OGG1 yielded the highest number of validated hits (18) and the most potent inhibitors (3). Several compounds, most notably GDC-0276, inhibited multiple but not all DNA glycosylases, suggesting the possibility of shared ligand-recognition features across subsets of the family, or a rarer assay interference mechanism.

**Figure 3.**
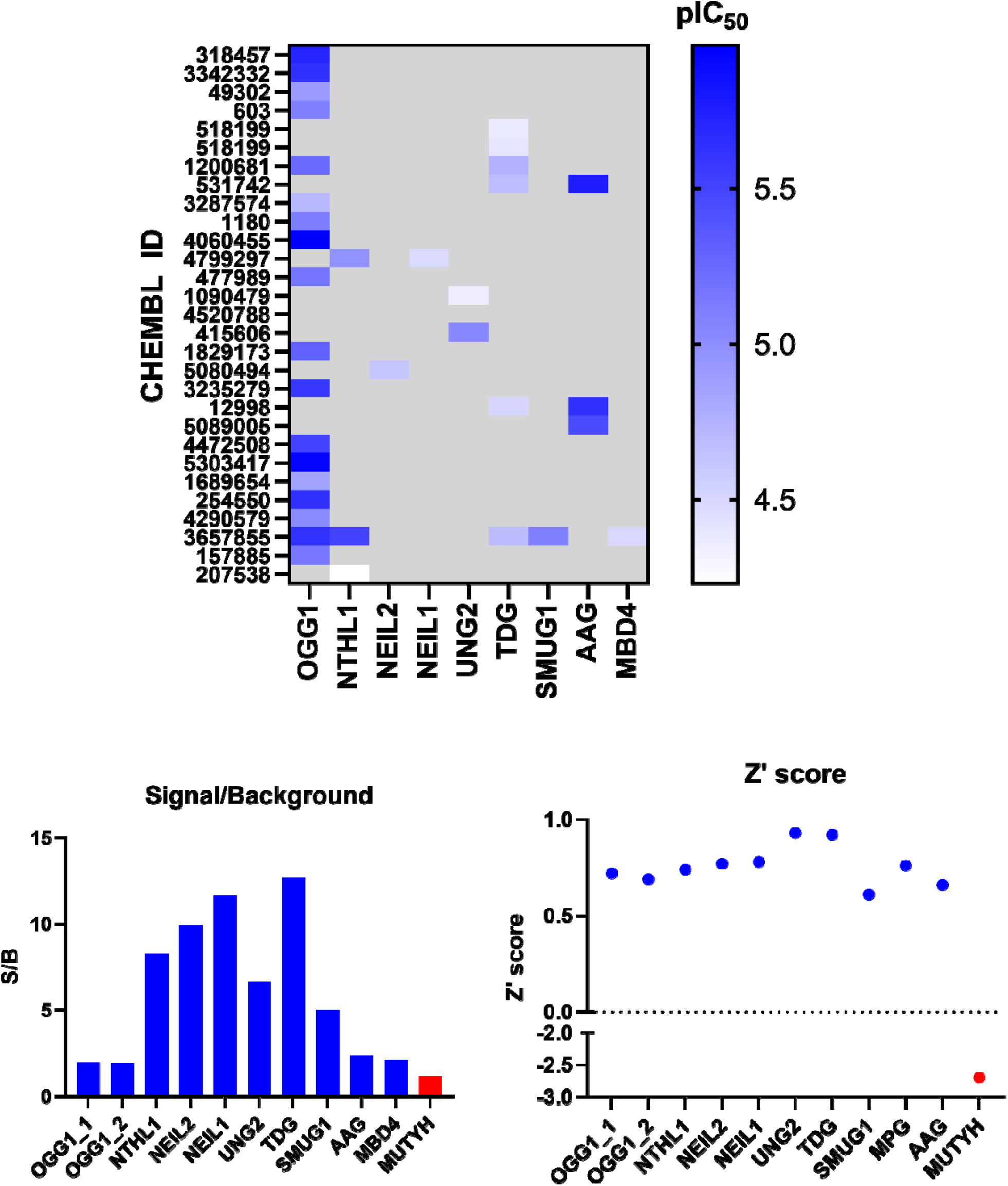
Hit validation: assay performance metrics across the glycosylase panel. Potencies (IC□□) of primary-screen hit compounds against corresponding enzymes. (Top). The quality and robustness of each DNA glycosylase assay **were** evaluated using the Z′-score (left) and signal-to-background (S/B) ratio (right). MUTYH (highlighted in red) showed reduced assay performance compared with the other DNA glycosylases. OGG1_1 and OGG1_2 denote two independent OGG1 validation plates, included separately because the number of validated OGG1 hits exceeded the capacity of a single assay plate.

**Figure 4.**
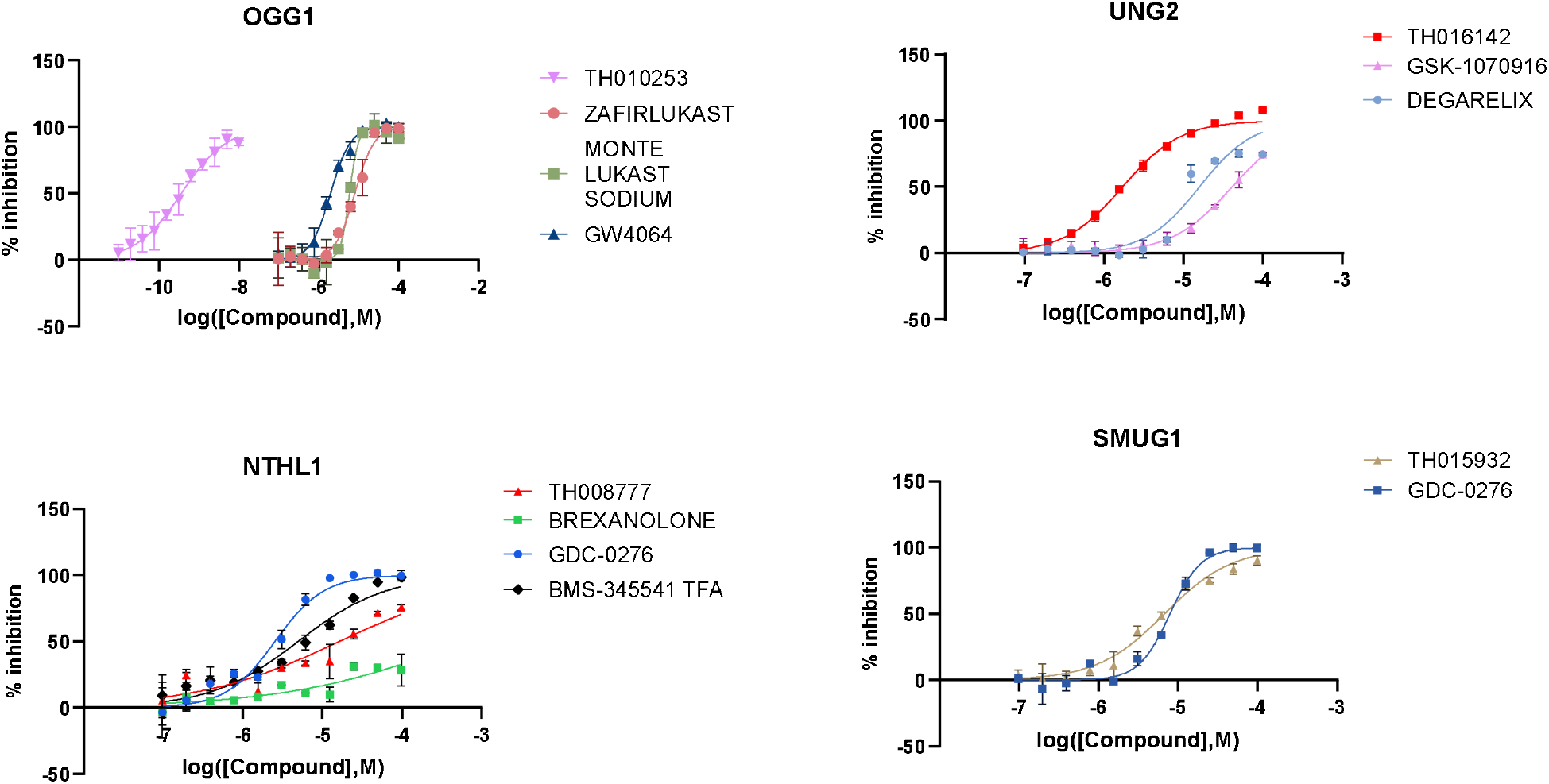
Concentration-response validation of selected DNA glycosylase inhibitors. Representative compounds identified in the primary screen were evaluated in concentration-response assays against OGG1, UNG2, NTHL1 and SMUG1. Enzyme activity is expressed as percent inhibition relative to assay controls and plotted as a function of compound concentration. Data were fitted using a four-parameter concentration-response model to determine inhibitory potency (pIC□□). Data points represent mean ± SD.

Using the positive and negative control wells on each assay plate, Z′ score and signal to background ratio (S/B) were calculated as a measure of assay performance (Figure 3). Assay performance was robust for nine glycosylases, with Z′ factors generally ranging from approximately 0.6 to 0.9. MUTYH and NEIL3 did not meet the predefined assay-quality criteria required for reliable hit identification and were therefore excluded from hit analysis. Otherwise, the results demonstrate that the assay conditions provided sufficient dynamic range and reproducibility for inhibitor screening across the DNA glycosylase family.

For OGG1, NTHL1 and UNG, selected validated hits were analysed by nanoDSF to further assess direct compound engagement with the target protein. A change in the melting temperature (T_M_) relative to the DMSO control is reported in Tables S1-S9. Selected compounds are shown in Table 2. For OGG1, the largest positive shifts were observed for zafirlukast (ΔTm = +7.54 °C), montelukast sodium (ΔTm = +5.41 °C), and GW4064 (ΔTm = +4.66 °C), consistent with direct target engagement. Interestingly, thermal responses were not uniformly stabilising. Several active compounds, including GDC-0276, GNE-131 and LY-223982, induced pronounced decreases in melting temperature despite measurable inhibitory activity. For UNG, two validated inhibitors were evaluated by nanoDSF. GSK-1070916 produced a positive thermal shift of +2.16 °C, consistent with direct protein engagement, whereas no thermal shift was observed for degarelix under the conditions tested. BMS-345541 TFA and brexanolone both induced negative thermal shifts (ΔTm = -2.09 °C and -1.56 °C, respectively) for NTHL1. Together, these findings indicate that enzymatic inhibition does not necessarily correlate with thermal stabilisation The nanoDSF dataset therefore provides an additional resource for prioritizing compounds for future chemical probe and medicinal chemistry efforts.

**Table 2.**
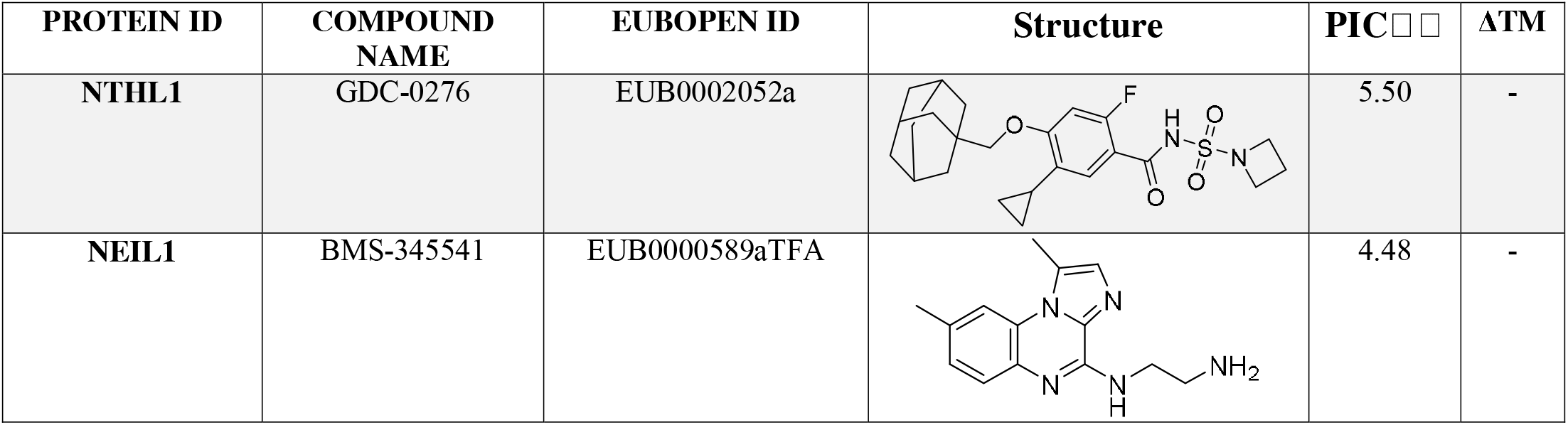
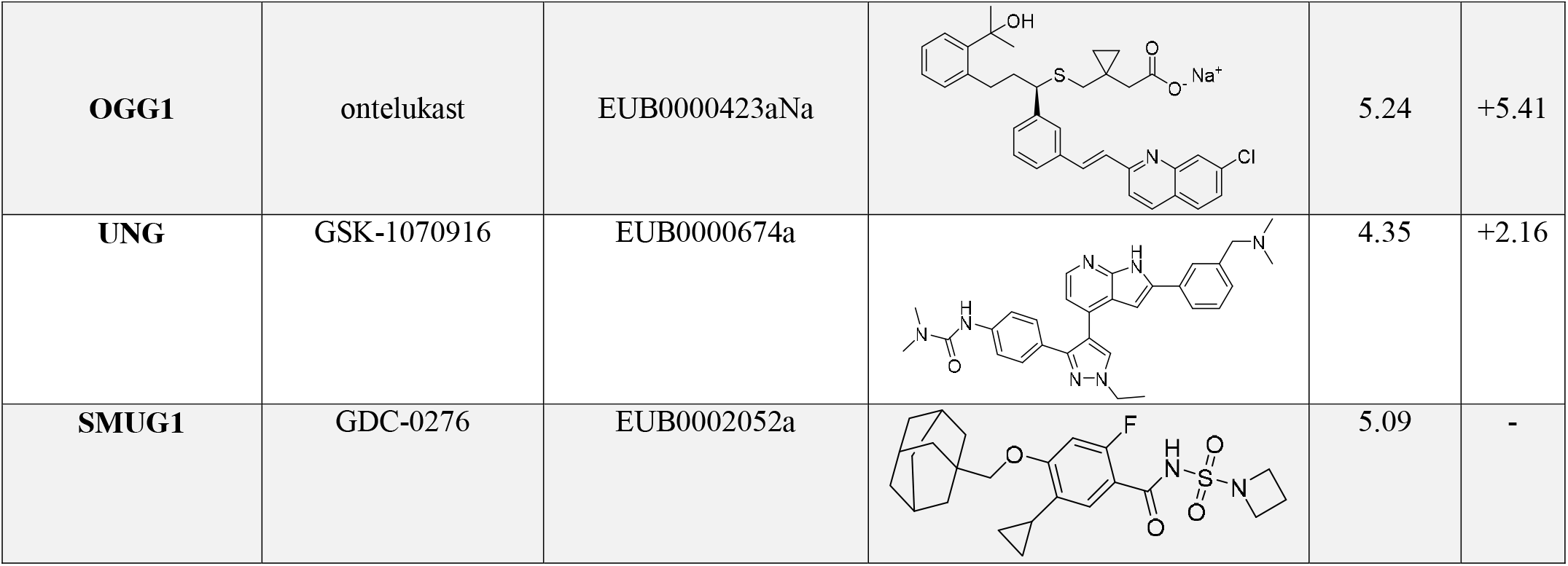
Selected compounds identified for selected DNA glycosylase targets. Protein targets, compound names, EUBOPEN identifiers, chemical structures, pIC□□ values, and thermal stabilization (ΔT□) values are shown.

| PROTEIN ID | COMPOUND NAME | EUBOPEN ID | Structure | $\text{pIC}_{50}$ | $\Delta T_m$ |
| --- | --- | --- | --- | --- | --- |
| NTHL1 | GDC-0276 | EUB0002052a |  | 5.50 | - |
| NEIL1 | BMS-345541 | EUB0000589aTFA |  | 4.48 | - |
| **OGG1** | ontelukast | EUB0000423aNa |  | 5.24 | +5.41 |
| **UNG** | GSK-1070916 | EUB0000674a |  | 4.35 | +2.16 |
| **SMUG1** | GDC-0276 | EUB0002052a |  | 5.09 | - |

### Data Availability

All primary screening data generated in this study have been deposited in ChEMBL and are publicly accessible through the assay identifiers listed in Table 3. The deposited records contain the compound-level activity data for the individual DNA glycosylase assays, enabling independent analysis and comparison across the enzyme panel. Together with the assay protocols and experimental information provided in this manuscript and its Supporting Information, these data are intended to provide a reusable resource for DNA glycosylase assay development, compound profiling and future chemical probe discovery.

**Table 3.** ChEMBL accession numbers for the DNA glycosylase screening datasets. ChEMBL assay identifiers corresponding to the primary small-molecule screening data generated for each DNA glycosylase are listed. The associated records provide access to the compound-level activity measurements reported in this study.

| TARGET | CHEMBL ASSAY ID |
| --- | --- |
| **MBD4** | CHEMBL5665444 |
| **AAG** | CHEMBL5665445 |
| **MUTYH** | CHEMBL5665446 |
| **NEIL1** | CHEMBL5665447 |
| **NEIL2** | CHEMBL5665448 |
| **NTHL1** | CHEMBL5665449 |
| **OGG1** | CHEMBL5665450 |
| **SMUG1** | CHEMBL5665451 |
| **TDG** | CHEMBL5665452 |
| **UNG** | CHEMBL5665453 |

## Discussion

DNA glycosylases initiate BER and collectively recognize a remarkably diverse spectrum of damaged and modified DNA bases. Despite their central role in genome maintenance, the chemical biology of this enzyme family remains unevenly developed. OGG1 has emerged as a tractable target for small-molecule modulation,^14,28-30^ and inhibitors have also been reported for selected other DNA glycosylases.^6-8,31^ However, for much of the family, chemical matter remains scarce and has generally been identified in studies focusing on individual enzymes rather than through systematic comparison across the glycosylase family. Here, we establish a common biochemical screening platform for human DNA glycosylases and use it to generate a family-wide dataset of small-molecule inhibition. The resulting resource substantially expands the chemical space associated with DNA glycosylases and, importantly, provides starting points for enzymes for which few or no suitable small-molecule inhibitors have previously been available.

The identification of inhibitors across structurally and functionally distinct DNA glycosylases further supports the concept that this protein family is amenable to small-molecule modulation.^8^ DNA-binding proteins have traditionally been considered challenging targets because their interaction surfaces are frequently large, dynamic and polar.^32^ We previously investigated this question computationally across DNA glycosylases and found potentially druggable pockets despite their low sequence conservation and the substantial conformational changes accompanying DNA recognition.^8^ The present experimental dataset extends this concept across the family. Although hit rates were generally modest, reproducible inhibition was observed for most glycosylases under screening conditions, with 37 of 63 compounds selected from the primary screen confirmed in concentration-response experiments. The relatively low hit rates combined with a high validation rate are encouraging characteristics for future screening campaigns and suggest that the observed activities are not simply a consequence of widespread nonspecific inhibition.

An important feature of the dataset is the considerable variation in chemical susceptibility between individual DNA glycosylases. OGG1 produced both the largest number of hits and some of the most potent compounds, consistent with its established tractability as a small-molecule target. In contrast, considerably fewer compounds were identified for several other members of the family. These differences should not necessarily be interpreted as intrinsic differences in protein druggability. The present screen samples a defined chemogenomic chemical space and the individual assays differ in substrate, enzyme concentration and, for monofunctional glycosylases, their requirement for APE1. Nevertheless, comparison of these profiles provides a first experimental attempt for examining how chemical recognition differs across the human DNA glycosylase family.

The cross-target activity observed for a subset of compounds is similarly informative. Several compounds inhibited more than one DNA glycosylase, with GDC-0276 representing a prominent example.^33^ Such activity may reflect shared physicochemical requirements for binding to glycosylase active sites or DNA-binding interfaces despite the limited sequence conservation across the family. Alternatively, activity across several assays can arise through mechanisms that are independent of direct glycosylase binding, including interactions with DNA, assay components or, in coupled assays, APE1. The inclusion of an APE1 counter-screen therefore provides an important level of annotation for the dataset. More generally, compounds displaying activity against multiple glycosylases should be regarded as useful starting points for mechanistic investigation rather than immediately classified as family-selective inhibitors. Conversely, differences in activity among related compounds in the CGL may provide useful preliminary structure-activity relationships from which greater potency and selectivity can be developed.

For selected inhibitors of OGG1, NTHL1 and UNG, nanoDSF provided an orthogonal assessment of compound protein interaction. Several inhibitors produced substantial changes in protein melting temperature, including pronounced thermal stabilisation of OGG1 by zafirlukast, montelukast and GW4064. Other active compounds instead decreased protein thermal stability, demonstrating that inhibitory activity does not necessarily translate into a positive thermal shift. Destabilisation can result from ligand-induced changes in protein conformation or conformational dynamics and should therefore not in itself be interpreted as absence of target engagement.^34^ At the same time, the lack of a thermal shift cannot exclude binding.^35^ Accordingly, the nanoDSF measurements are best considered complementary to the biochemical inhibition data and provide an additional parameter for prioritising compounds for mechanistic, structural and medicinal chemistry studies.

The identification of activity among compounds originally developed for other protein targets also raises a second potential use of this dataset. Chemogenomic libraries deliberately contain pharmacologically annotated compounds and therefore provide an opportunity to identify previously unrecognised target activities.^36^ Such interactions may contribute to cellular phenotypes attributed to these molecules or reveal connections between DNA repair and other biological pathways. Zafirlukast is an interesting example in this respect. The compound, a leukotriene receptor antagonist which is clinically used for chronic treatment and prevention of asthma and, in the present dataset, inhibits OGG1 and induces a substantial positive thermal shift. This observation is notable given the increasingly recognised functions of OGG1 beyond canonical repair, particularly in the regulation of inflammatory gene expression.^14,37^ Genetic depletion of OGG1 reduces allergic airway inflammation, and pharmacological inhibition of OGG1 has similarly been shown to suppress inflammatory responses and airway hyperresponsiveness in experimental models.^38,39^ Thus, while the biochemical activity identified here does not establish that OGG1 contributes to the pharmacological effects of zafirlukast, the intersection provides a testable hypothesis and illustrates how systematic profiling datasets can reveal unexpected connections between established pharmacology and DNA repair biology.

The compounds identified here should primarily be considered starting points rather than immediately applicable chemical probes. Most inhibitors display micromolar potency, and comprehensive selectivity profiling, structural characterisation and cellular target-engagement studies will be required before individual molecules can be used to assign biological functions to a particular glycosylase. This distinction is especially important for DNA glycosylases because compounds may influence enzymatic activity through several mechanisms, including direct occupation of the catalytic pocket, interference with DNA binding, interaction with the DNA substrate or modulation of individual catalytic steps. Indeed, work on OGG1 has demonstrated that small molecules interacting with DNA glycosylases can produce considerably more complex functional consequences than simple inhibition.^28^ The dataset presented here should therefore be viewed as the beginning of chemical probe development.

Beyond the individual compounds, a major outcome of this work is the establishment of a transferable screening platform for the human DNA glycosylase family.^12^ The use of related fluorophore-quencher substrates and a common screening format enables compounds to be compared across enzymes and facilitates rapid assessment of selectivity within the family. The assays may consequently be useful not only for reproducing the present screen but also for evaluating new chemical series, profiling compounds identified by orthogonal approaches and supporting medicinal chemistry optimisation. Together with the primary screening measurements, concentration-response data, assay-quality parameters, counter-screening and biophysical measurements, these protocols provide a reference dataset against which future DNA glycosylase inhibitors can be evaluated.

Making these data openly available is particularly important for targets for which chemical tools remain underdeveloped. Large, consistently generated protein-ligand datasets are increasingly valuable not only for conventional chemical biology but also for computational approaches to ligand prediction and machine learning. Here, the combination of active and inactive compounds across related enzymes is potentially as informative as the individual hits, as it captures target-specific and family-wide patterns of chemical recognition. Deposition of the complete dataset, including negative results and experimental metadata, should therefore enable uses beyond those anticipated in the present study and allow the data to be reanalysed as new computational and experimental approaches emerge.

In summary, we provide a systematic small-molecule screening dataset across the human DNA glycosylase family and identify validated inhibitors for multiple members of this important class of DNA repair enzymes. The work expands the available chemical matter for DNA glycosylases, establishes compounds that can serve as assay controls and starting points for chemical probe development, and provides a common experimental framework for profiling future inhibitors. Most importantly, by making the complete screening data and protocols openly available, this resource lowers the experimental barrier to studying chemical modulation of DNA glycosylases and provides a foundation for the development of more potent, selective and mechanistically defined tools to interrogate their functions in genome maintenance and disease.

## AUTHOR INFORMATION

### Corresponding Author

Authorship has been determined in accordance with the ICMJE (Vancouver) authorship guidelines. CRediT taxonomy was used to determine order where possible. Remaining authors are displayed alphabetically.

## Acknowledgements

We thank the taxpayers of our individual countries for their generous support of our research. We thank the scientists of the EUbOPEN consortium and those at SciLifeLab, CBCS, CMM and Biomedicum for their support and access to infrastructure. We would like to thank Athina Pliakou, Mari Kullman Magnusson and Kristina Edfeldt.

## References

1 David, S. S., O’Shea, V. L. & Kundu, S. Base-excision repair of oxidative DNA damage. Nature 447, 941–950 (2007). 10.1038/nature05978

2 Visnes, T. et al. Targeting BER enzymes in cancer therapy. DNA Repair (Amst) 71, 118–126 (2018). 10.1016/j.dnarep.2018.08.015

3 Hans, F., Senarisoy, M., Bhaskar Naidu, C. & Timmins, J. Focus on DNA Glycosylases-A Set of Tightly Regulated Enzymes with a High Potential as Anticancer Drug Targets. Int J Mol Sci 21 (2020). 10.3390/ijms21239226

4 Mechetin, G. V., Endutkin, A. V., Diatlova, E. A. & Zharkov, D. O. Inhibitors of DNA Glycosylases as Prospective Drugs. Int J Mol Sci 21 (2020). 10.3390/ijms21093118

5 Renaudin, X. & Campalans, A. Modulation of OGG1 enzymatic activities by small molecules, promising tools and current challenges. DNA Repair (Amst) 149, 103827 (2025). 10.1016/j.dnarep.2025.103827

6 Zhou, J.-X. et al. Targeting thymine DNA glycosylase induces synthetic lethality in p53-deficient cancers. Nature Chemical Biology 22, 1109–1119 (2026). 10.1038/s41589-025-02100-1

7 Thacker, P. S. et al. A SMUG1 Inhibitor Modulates the Excision of Pyrimidine DNA Damage. ACS Medicinal Chemistry Letters 17, 1285–1293 (2026). 10.1021/acsmedchemlett.6c00051

8 Michel, M. et al. Computational and Experimental Druggability Assessment of Human DNA Glycosylases. ACS Omega 4, 11642–11656 (2019). 10.1021/acsomega.9b00162

9 Ackloo, S. et al. Target 2035 - an update on private sector contributions. RSC Med Chem 14, 1002–1011 (2023). 10.1039/d2md00441k

10 Müller, S. et al. Target 2035 - update on the quest for a probe for every protein. RSC Med Chem 13, 13–21 (2022). 10.1039/d1md00228g

11 Edwards, A. M. et al. Protein–ligand data at scale to support machine learning. Nature Reviews Chemistry 9, 634–645 (2025). 10.1038/s41570-025-00737-z

12 Protocol for DNA Glycolysases, < https://www.eubopen.org/protocols-reagents > (2021).

13 Wallner, O. et al. Optimization of N-Piperidinyl-Benzimidazolone Derivatives as Potent and Selective Inhibitors of 8-Oxo-Guanine DNA Glycosylase 1. ChemMedChem 18, e202200310 (2023). 10.1002/cmdc.202200310

14 Visnes, T. et al. Small-molecule inhibitor of OGG1 suppresses proinflammatory gene expression and inflammation. Science 362, 834–839 (2018). 10.1126/science.aar8048

15 Haslam, J. et al. A monofunctional-like mutant of DNA glycosylase NTHL1 changes the dynamics of DNA repair during acute oxidative stress. J Biol Chem 302, 111332 (2026). 10.1016/j.jbc.2026.111332

16 O’Brie, P. J. & Ellenberger, T. Human Alkyladenine DNA Glycosylase Uses Acid−Base Catalysis for Selective Excision of Damaged Purines. Biochemistry 42, 12418–12429 (2003). 10.1021/bi035177v

17 Krokeide, S. Z. et al. Human NEIL3 is mainly a monofunctional DNA glycosylase removing spiroimindiohydantoin and guanidinohydantoin. DNA Repair (Amst) 12, 1159–1164 (2013). 10.1016/j.dnarep.2013.04.026

18 Hank, E. C. et al. Nucleobase catalysts for the enzymatic activation of 8-oxoguanine DNA glycosylase 1. RSC Chem Biol 7, 169–181 (2026). 10.1039/d4cb00323c

19 Tredup, C. et al. Toward target 2035: EUbOPEN - a public-private partnership to enable & unlock biology in the open. RSC Med Chem 16, 457–464 (2025). 10.1039/d4md00735b

20 EUbOPEN Gateway, < https://gateway.eubopen.org/prototypes/sets > (

21 Harding, S. D. et al. The IUPHAR/BPS Guide to PHARMACOLOGY in 2024. Nucleic Acids Res 52, D1438–d1449 (2024). 10.1093/nar/gkad944

22 Harding, S. D. et al. The IUPHAR/BPS Guide to PHARMACOLOGY in 2026. Nucleic Acids Research 54, D1446–D1456 (2026). 10.1093/nar/gkaf1067

23 Škuta, C., Southan, C. & Bartůněk, P. Will the chemical probes please stand up? RSC Medicinal Chemistry 12, 1428–1441 (2021). 10.1039/d1md00138h

24 Skuta, C. et al. Probes & Drugs portal: an interactive, open data resource for chemical biology. Nature Methods 14, 759–760 (2017). 10.1038/nmeth.4365

25 Zdrazil, B. et al. The ChEMBL Database in 2023: a drug discovery platform spanning multiple bioactivity data types and time periods. Nucleic Acids Research 52, D1180–D1192 (2024). 10.1093/nar/gkad1004

26 Hank, E. C. et al. Nucleobase catalysts for the enzymatic activation of 8-oxoguanine DNA glycosylase 1. RSC Chem Biol (2025). 10.1039/d4cb00323c

27 Varga, M. et al. Giving an Enzyme Scissors: Serotonin Derivatives as Potent Organocatalytic Switches for DNA Repair Enzyme OGG1. Journal of Medicinal Chemistry (2025). 10.1021/acs.jmedchem.5c01454

28 Michel, M. et al. Small-molecule activation of OGG1 increases oxidative DNA damage repair by gaining a new function. Science 376, 1471–1476 (2022). 10.1126/science.abf8980

29 Tahara, Y.-k. et al. Potent and Selective Inhibitors of 8_Oxoguanine DNA Glycosylase. Journal of the American Chemical Society 140, 2105–2114 (2018). 10.1021/jacs.7b09316

30 Donley, N. et al. Small Molecule Inhibitors of 8_Oxoguanine DNA Glycosylase_1 (OGG1). ACS Chemical Biology 10, 2334–2343 (2015). 10.1021/acschembio.5b00452

31 Gao, Y. et al. Small-molecule activator of SMUG1 enhances repair of pyrimidine lesions in DNA. DNA Repair (Amst) 146, 103809 (2025). 10.1016/j.dnarep.2025.103809

32 Srinivasan, A. & Gold, B. Small-molecule inhibitors of DNA damage-repair pathways: an approach to overcome tumor resistance to alkylating anticancer drugs. Future Med Chem 4, 1093–1111 (2012). 10.4155/fmc.12.58

33 Rothenberg, M. E. et al. Safety, Tolerability, and Pharmacokinetics of GDC-0276, a Novel Na(V)1.7 Inhibitor, in a First-in-Human, Single- and Multiple-Dose Study in Healthy Volunteers. Clin Drug Investig 39, 873–887 (2019). 10.1007/s40261-019-00807-3

34 Gao, K., Oerlemans, R. & Groves, M. R. Theory and applications of differential scanning fluorimetry in early-stage drug discovery. Biophys Rev 12, 85–104 (2020). 10.1007/s12551-020-00619-2

35 Rombouts, F. J. R. et al. Fragment Binding to β-Secretase 1 without Catalytic Aspartate Interactions Identified via Orthogonal Screening Approaches. ACS Omega 2, 685–697 (2017). 10.1021/acsomega.6b00482

36 Schoppa, M. Q. & Drewry, D. H. Chemogenomic sets: valuable tools for early-stage drug discovery. Essays Biochem (2026). 10.1042/ebc20260007

37 Visnes, T. et al. Targeting OGG1 arrests cancer cell proliferation by inducing replication stress. Nucleic Acids Res 48, 12234–12251 (2020). 10.1093/nar/gkaa1048

38 Tanner, L. et al. Small-molecule-mediated OGG1 inhibition attenuates pulmonary inflammation and lung fibrosis in a murine lung fibrosis model. Nat Commun 14, 643 (2023). 10.1038/s41467-023-36314-5

39 Li, G. et al. 8-Oxoguanine-DNA glycosylase 1 deficiency modifies allergic airway inflammation by regulating STAT6 and IL-4 in cells and in mice. Free Radic Biol Med 52, 392–401 (2012). 10.1016/j.freeradbiomed.2011.10.490

