## Supporting Information for "A high-throughput screening dataset of small-molecule inhibitors across human DNA glycosylases"

**Figure S1. Primary screening of the CGL against human DNA glycosylases MBD4, NTHL1, AAG/MPG, TDG, SMUG1 and NEIL2.** Compounds were screened against the indicated DNA glycosylases and activity is shown as percent inhibition relative to plate controls. Each blue point represents an individual compound and red stars indicate compounds selected as primary hits. Red dashed lines indicate the plate-specific thresholds used for hit selection, and vertical dashed lines separate individual screening plates. Compound numbers correspond to their position within the screening library. Remaining members are shown in Figure 2.

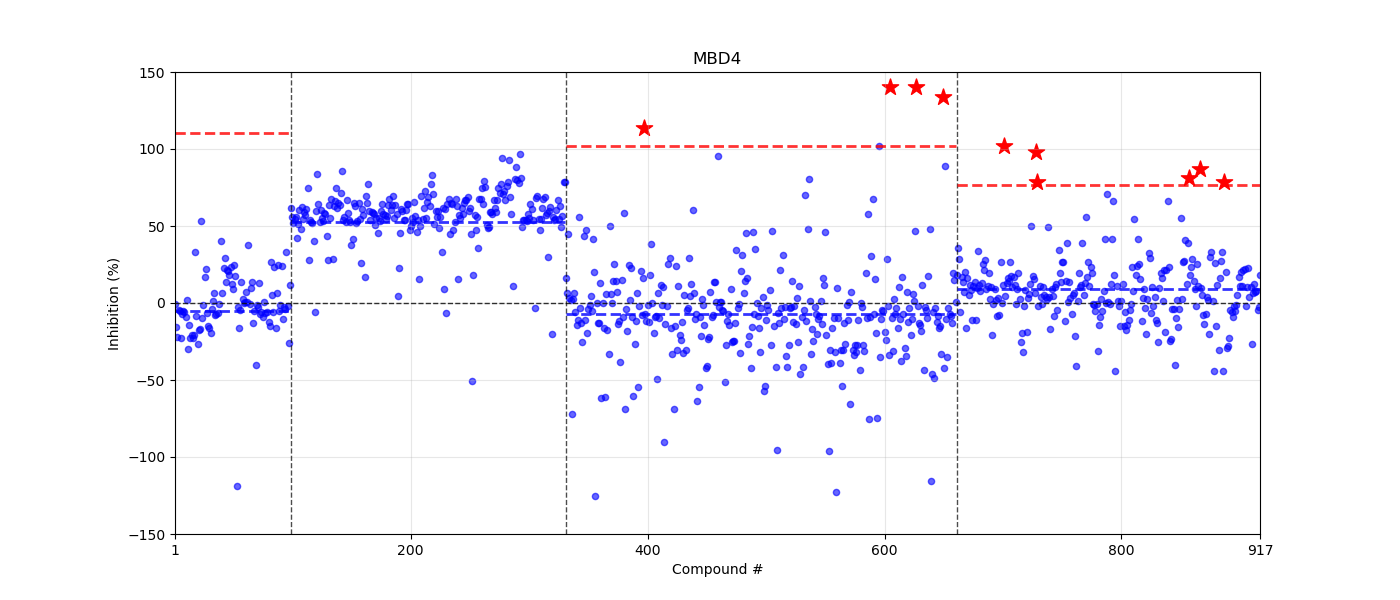

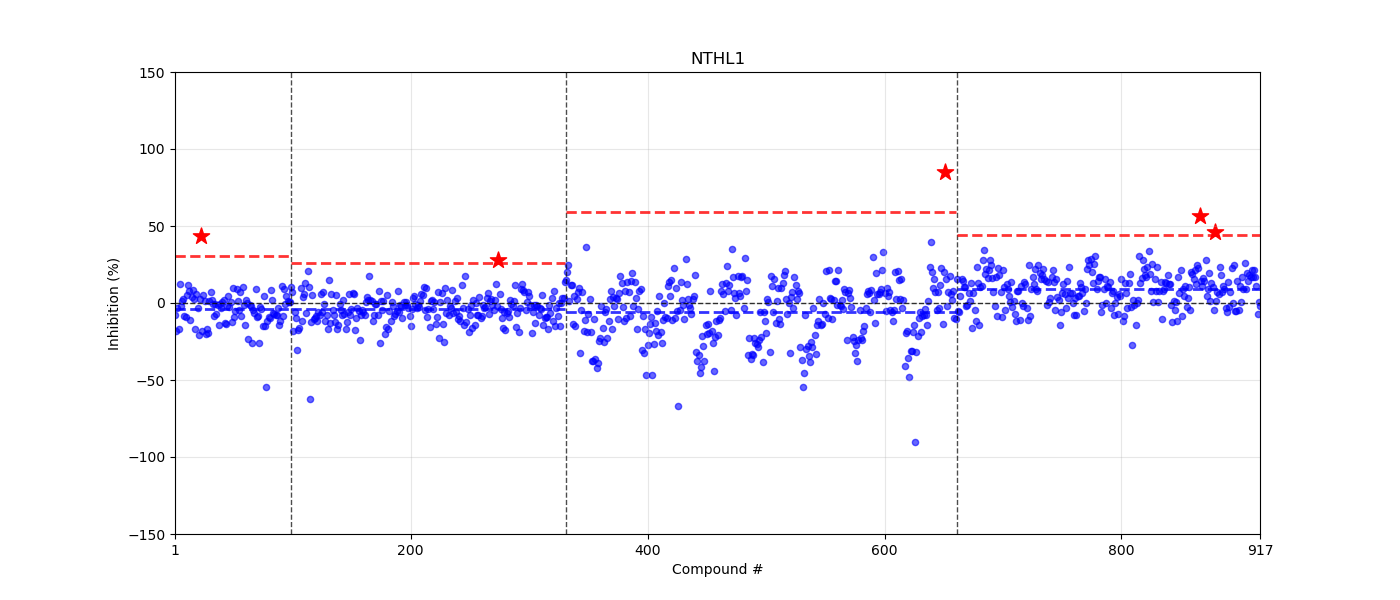

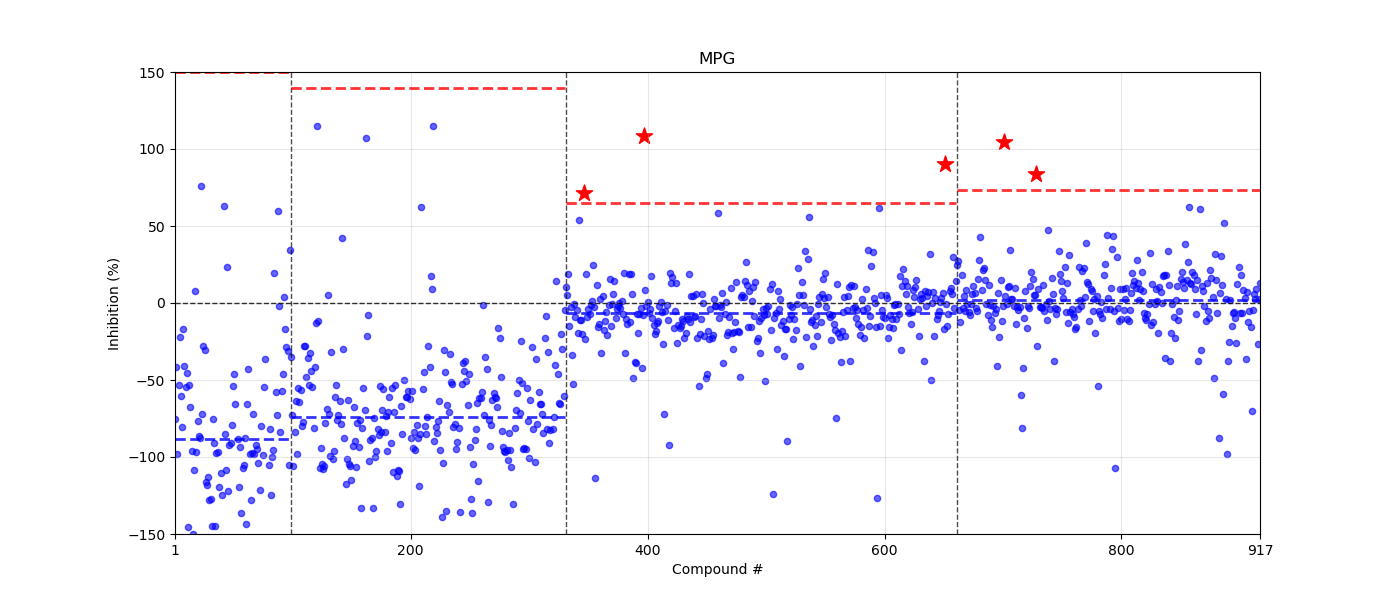

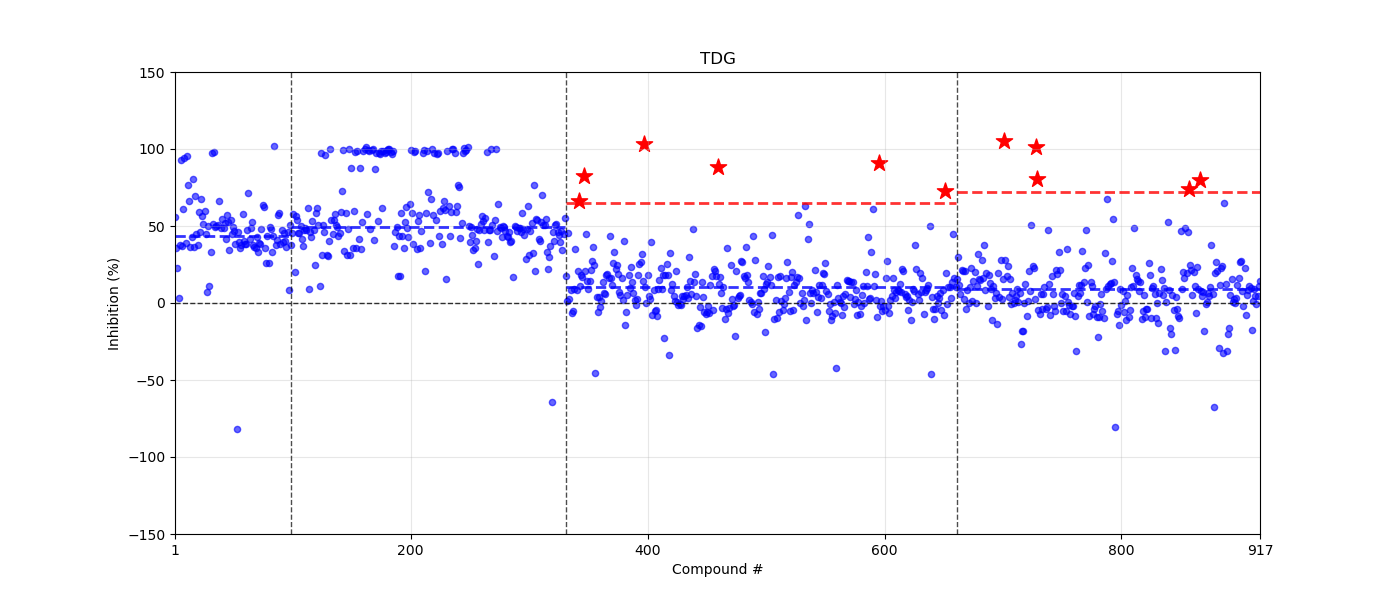

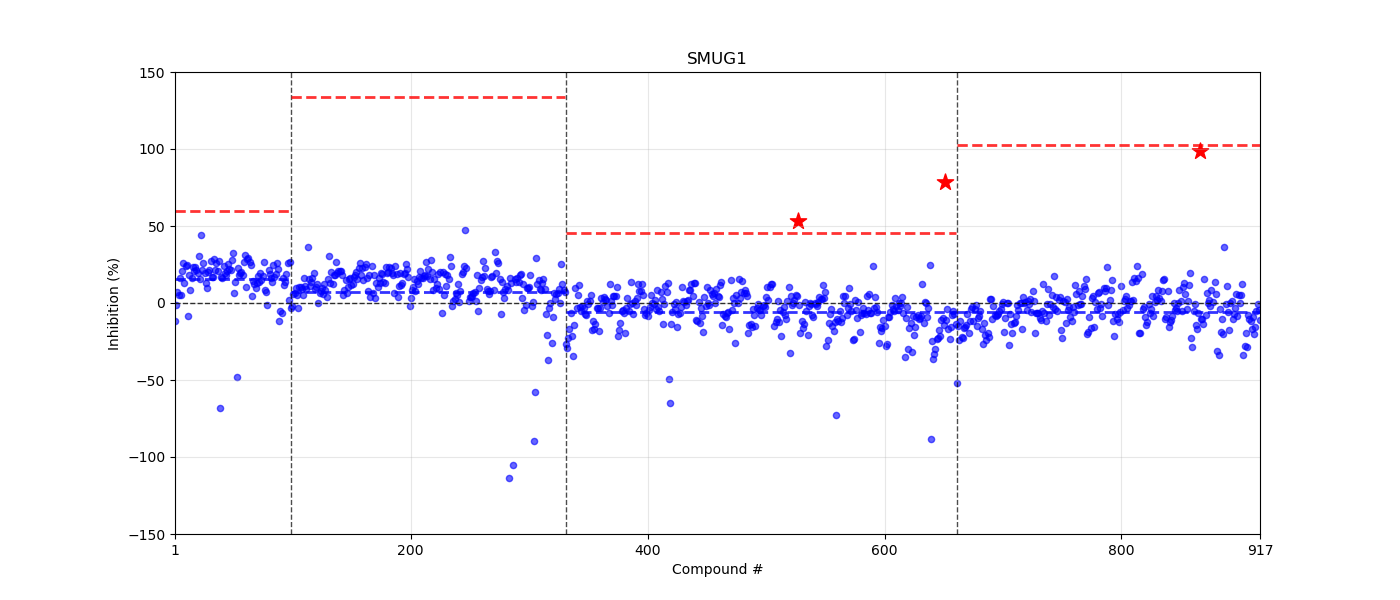

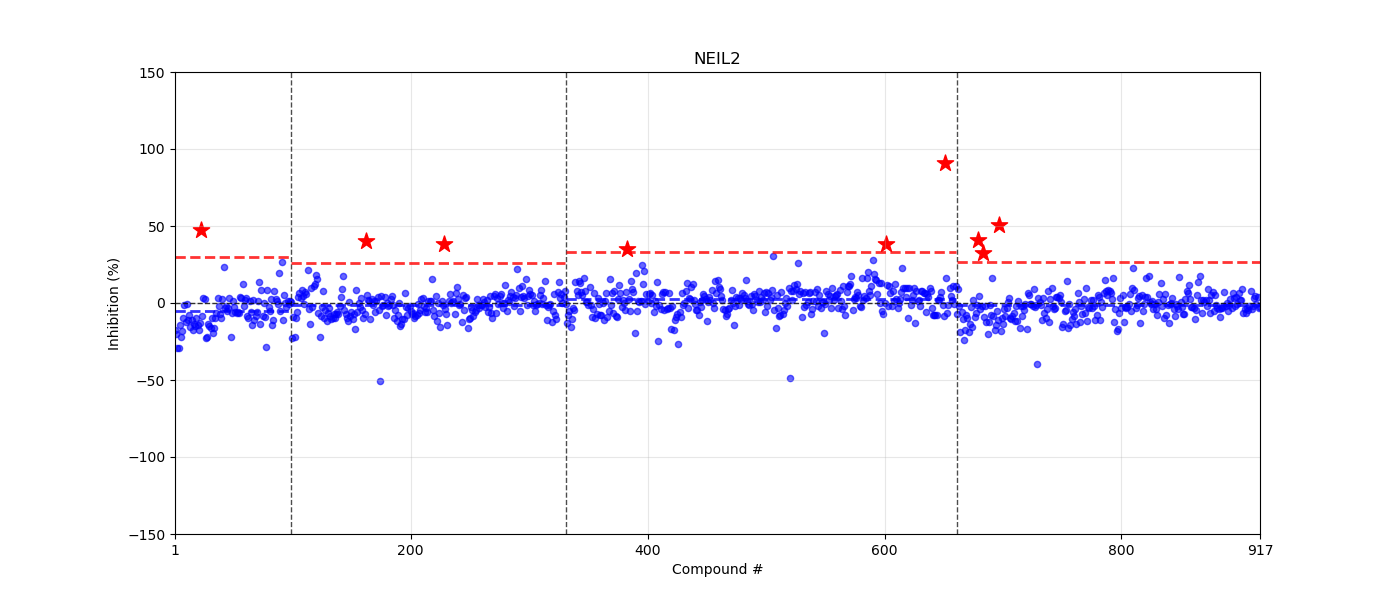

**Table S1. Confirmed inhibitors of OGG1, ordered by pIC150**

| Compound Name | Compound ChEMBL ID | Compound ID | Structure | pIC_50_ | ΔT_m ,_ °C |
| --- | --- | --- | --- | --- | --- |
| GW3965 HCl | CHEMBL4060455 | EUB0000572aCl | 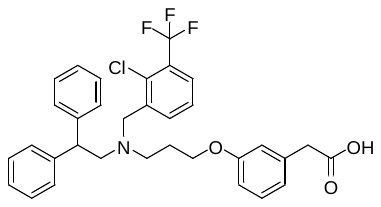 | 5.96 | -2.61 |
| RGX-104 HCl | CHEMBL5303417 | EUB0001482aCl | 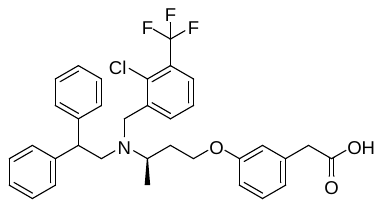 | 5.93 | -1.73 |
| GW4064 | CHEMBL318457 | EUB0000184a | 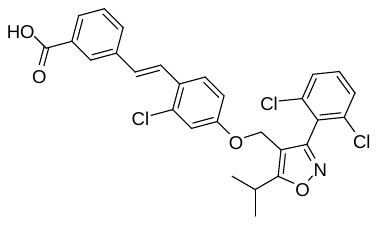 | 5.71 | +4.66 |
| A-1155463 | CHEMBL3342332 | EUB0000262a | 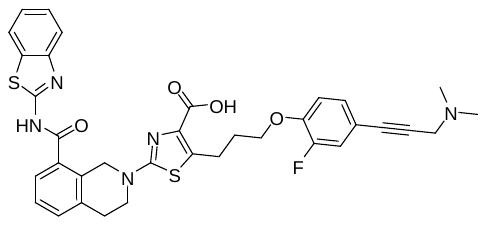 | 5.63 | +1.55 |
| polygodial | CHEMBL254550 | EUB0001996a | 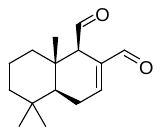 | 5.63 | - |
| GDC-0276 | CHEMBL3657855 | EUB0002052a | 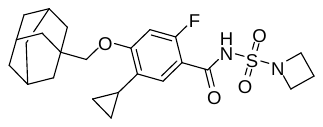 | 5.62 | -9.75 |
| capadenoson | CHEMBL3235279 | EUB0000948a | 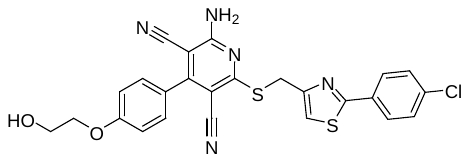 | 5.58 | +1.42 |
| cintirorgon | CHEMBL4472508 | EUB0001172a | 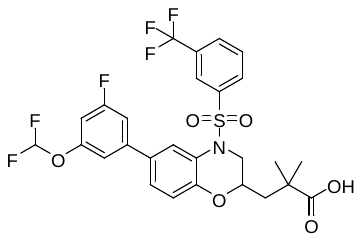 | 5.51 | -4.8 |
| AMG-837 | CHEMBL1829173 | EUB0000855a | 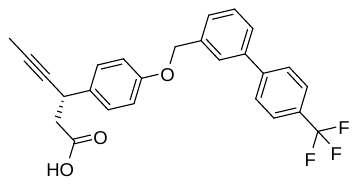 | 5.3 | -2.74 |
| montelukast sodium | CHEMBL1200681 | EUB0000423aNa | 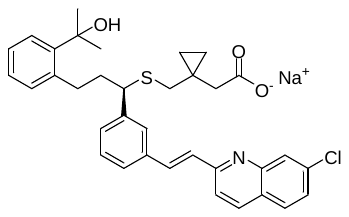 | 5.24 | +5.41 |
| SB590885 | CHEMBL477989 | EUB0000655a | - | 5.17 | +0.67 |
| KB-130015 | CHEMBL157885 | EUB0002355a | 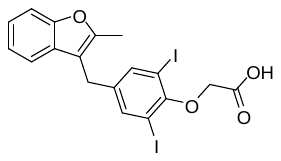 | 5.15 | -0.73 |
| AHPN | CHEMBL1180 | EUB0000568a | 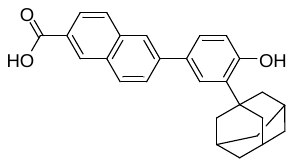 | 5.1 | +1.57 |
| zafirlukast | CHEMBL603 | EUB0000407a | 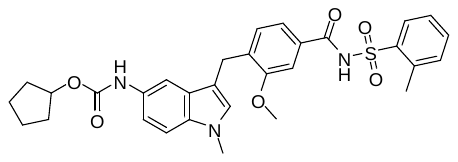 | 5.08 | +7.54 |
| GNE-131 | CHEMBL4290579 | EUB0002043a | 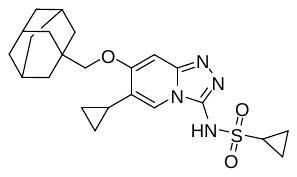 | 5.02 | -9.36 |
| LY-223982 | CHEMBL49302 | EUB0000370a | 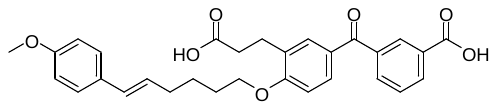 | 4.92 | -9.44 |
| QO 58 | CHEMBL1689654 | EUB0001978a | 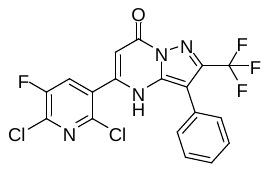 | 4.86 | -0.13 |
| AM-4668 | CHEMBL3287574 | EUB0000517a | 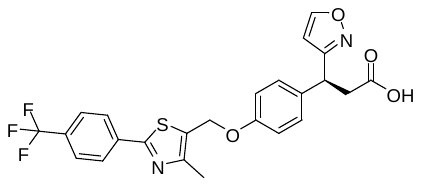 | 4.7 | -4.66 |

**Table S2. Confirmed inhibitors of NTHL1**

| Compound Name | Compound ChEMBL ID | Compound ID | | Structure | | pIC_50_ | | ΔT_m ,_ °C |
| --- | --- | --- | --- | --- | --- | --- | --- | --- |
| GDC-0276 | CHEMBL3657855 | EUB0002052a | 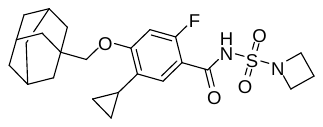 | | 5.50 | | - | |
| BMS-345541 TFA | CHEMBL4799297 | EUB0000589aTFA | 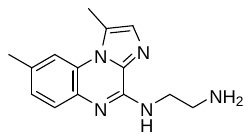 | | 4.97 | | -2.09 | |
| brexanolone | CHEMBL207538 | EUB0002367a | 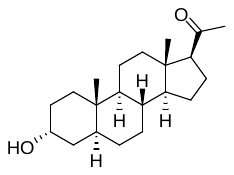 | | 4.23 | | -1.56 | |

**Table S3. NEIL1 inhibitor**

| Compound Name | Compound ChEMBL ID | Compound ID | Structure | pIC_50_ | ΔT_m ,_ °C |
| --- | --- | --- | --- | --- | --- |
| BMS-345541 TFA | CHEMBL4799297 | EUB0000589aTFA | 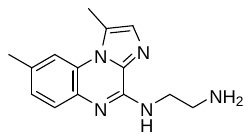 | 4.48 | - |

**Table S4. NEIL2 inhibitor**

| Compound Name | Compound ChEMBL ID | Compound ID | Structure | pIC_50_ | ΔT_m ,_ °C |
| --- | --- | --- | --- | --- | --- |
| L-803,087 TFA | CHEMBL5080494 | EUB0000882aTFA | 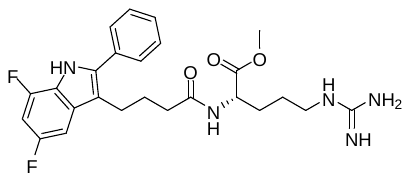 | 4.61 | - |

**Table S5. MBD4 inhibitor**

| Compound Name | Compound ChEMBL ID | Compound ID | Structure | pIC_50_ | ΔT_m ,_ °C |
| --- | --- | --- | --- | --- | --- |
| GDC-0276 | CHEMBL3657855 | EUB0002052a | 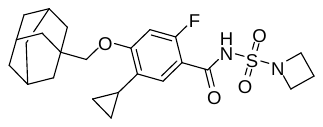 | 4.49 | - |

**Table S6. TDG inhibitors**

| Compound Name | Compound ChEMBL ID | Compound ID | | Structure | pIC_50_ | | ΔT_m ,_ °C |
| --- | --- | --- | --- | --- | --- | --- | --- |
| montelukast sodium | CHEMBL1200681 | EUB0000423aNa | | 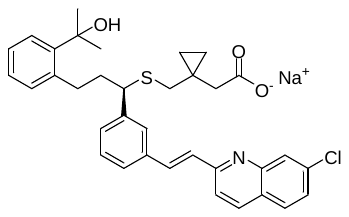 | 4.75 | |  |
| GDC-0276 | CHEMBL3657855 | EUB0002052a | | 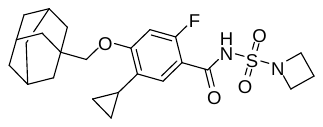 | 4.69 | |  |
| JTC-801 HCl | CHEMBL531742 | EUB0000427aCl | | 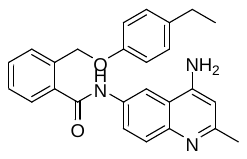 | 4.68 | |  |
| L-755507 | CHEMBL12998 | EUB0000985a | | 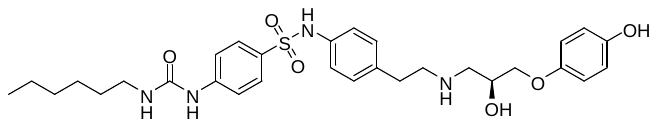 | 4.51 | |  |
| L-817,818 | CHEMBL518199 | EUB0000411b | 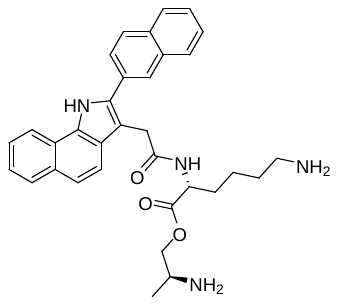 | | | 4.40 |  |
| L-817,818 | CHEMBL518199 | EUB0000411a | | 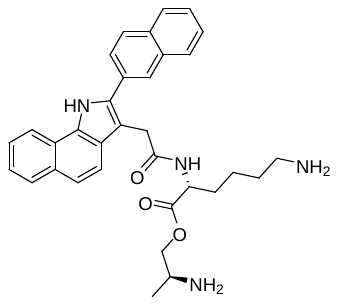 | 4.37 | |  |

**Table S7. UNG inhibitors**

| Compound Name | Compound ChEMBL ID | Compound ID | Structure | pIC_50_ | ΔT_m ,_ °C |
| --- | --- | --- | --- | --- | --- |
| degarelix | CHEMBL415606 | EUB0000760a | 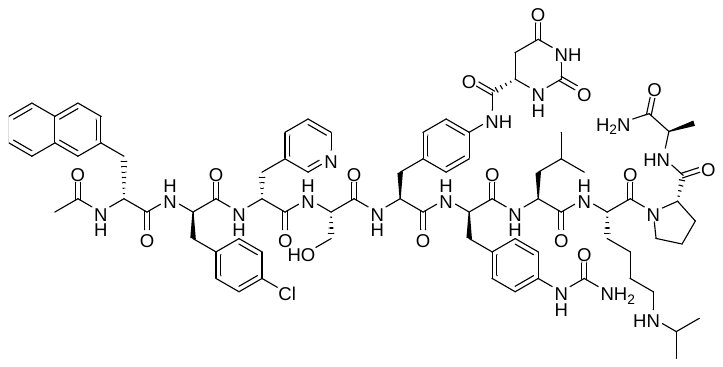 | 5.04 | - |
| GSK-1070916 | CHEMBL1090479 | EUB0000674a | 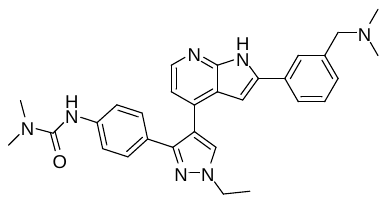 | 4.35 | +2.16 |

**Table S8. SMUG1 inhibitor**

| Compound Name | Compound ChEMBL ID | Compound ID | Structure | pIC_50_ | ΔT_m ,_ °C |
| --- | --- | --- | --- | --- | --- |
| GDC-0276 | CHEMBL3657855 | EUB0002052a | 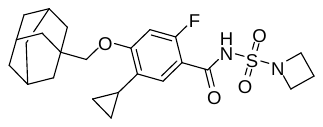 | 5.09 | - |

**Table S9. MPG inhibitors**

| Compound Name | Compound ChEMBL ID | Compound ID | Structure | pIC_50_ | ΔT_m ,_ °C |
| --- | --- | --- | --- | --- | --- |
| JTC-801 HCl | CHEMBL531742 | EUB0000427aCl | 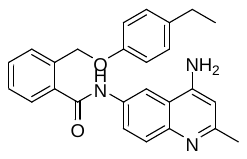 | 5.77 |  |
| L-755507 | CHEMBL12998 | EUB0000985a | 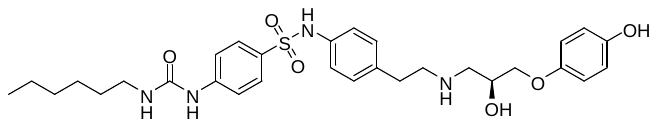 | 5.63 |  |
| SB 216641 HCl | CHEMBL5089005 | EUB0001016aCl | 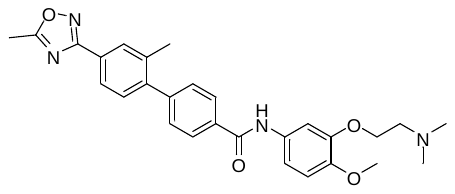 | 5.45 |  |

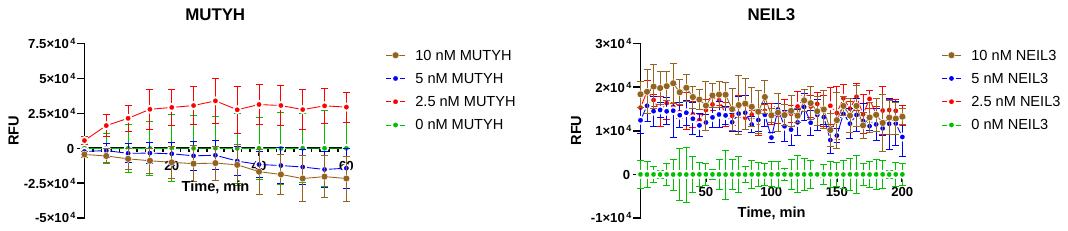

**Figure S1. Example enzyme titration plots for MUTYH and NEIL3 that were not carried forward for screening.** Fluorescence signals were monitored over time in the presence of increasing concentrations of MUTYH (0, 2.5, 5, and 10 nM; left) or NEIL3 (0, 2.5, 5, and 10 ng; right). Under the assay conditions tested, neither enzyme produced a sufficiently robust, concentration-dependent response to support implementation in the screening workflow. These assays were therefore not progressed to subsequent screening experiments.
